# Determining distinct roles of IL-1α through generation of an IL-1α knockout mouse with no defect in IL-1β expression

**DOI:** 10.1101/2022.09.21.508892

**Authors:** R.K. Subbarao Malireddi, Ratnakar Bynigeri, Balabhaskararao Kancharana, Bhesh Raj Sharma, Amanda R. Burton, Stephane Pelletier, Thirumala-Devi Kanneganti

## Abstract

Interleukin 1α (IL-1α) and IL-1β are the founding members of the IL-1 cytokine family, and these innate immune inflammatory mediators are critically important in health and disease. Early studies on these molecules suggested that their expression was interdependent, with an initial genetic model of IL-1α depletion, the IL-1α KO mouse (*Il1a*-KO^line1^), showing reduced IL-1β expression. However, studies using this line in models of infection and inflammation resulted in contrasting observations. To overcome the limitations of this genetic model, we have generated and characterized a new line of IL-1α KO mice (*Il1a*-KO^line2^) using CRISPR-Cas9 technology. In contrast to cells from the *Il1a*-KO^line1^, where IL-1β expression was drastically reduced, bone marrow-derived macrophages (BMDMs) from *Il1a*-KO^line2^ mice showed normal induction and activation of IL-1β. Additionally, *Il1a*-KO^line2^ BMDMs showed normal inflammasome activation and IL-1β expression in response to multiple innate immune triggers, including both pathogen-associated molecular patterns and pathogens. Moreover, using *Il1a*-KO^line2^ cells, we confirmed that IL-1α, independent of IL-1β, is critical for the expression of the neutrophil chemoattractant KC/CXCL1. Overall, we report the generation of a new line of IL-1α KO mice and confirm functions for IL-1α independent of IL-1β. Future studies on the unique functions of IL-1α and IL-1β using these mice will be critical to identify new roles for these molecules in health and disease and develop therapeutic strategies.

## INTRODUCTION

The IL-1 family of cytokines is a diverse family made up of potent inducers of inflammation. Members of this family can either prevent or promote disease, and they have been widely recognized as potential therapeutic targets (Lukens et al, 2012; Malik & Kanneganti, 2018; Ridker et al, 2017a; Ridker et al, 2017b; Ridker et al, 2011). The three members of the IL-1 sub-family, IL-1α, IL-1β, and IL-1 receptor antagonist (IL-1RA), bind the same IL-1 receptor (IL-1R). The cytokines IL-1α and IL-1β act as agonistic ligands, whereas IL-1RA is a strong antagonist; together, these molecules orchestrate robust proinflammatory immune responses (Dinarello, 2009; Dinarello et al, 1974).

Among the IL-1 cytokines, significant overlap has been observed in the downstream processes they activate. However, there are also key differences between their expression and release and the biological processes they drive (Cavalli et al, 2021). The pro-form of IL-1β is biologically inactive and requires proteolytic processing for its activation. Inflammasome-dependent caspase-1 activation and pyroptosis are the major mechanisms responsible for IL-1β processing and release (Kanneganti, 2010; Kayagaki et al, 2015; Shi et al, 2015). Unlike IL-1β, the pro-form of IL-1α is constitutively expressed in most cells from healthy hosts (Berda-Haddad et al, 2011; Kupper et al, 1986); it is also biologically active and can be present directly on the plasma membrane for signaling or released following membrane damage during various forms of cell death, making it a classic danger signal (Kaplanski et al, 1994; Kurt-Jones et al, 1985; Malik & Kanneganti, 2018).

As signaling molecules, a wide range of pathogen-associated and damage-associated molecular patterns (PAMPs and DAMPs) that activate innate immune signaling induce the expression and activation of both IL-1α and IL-1β (Malik & Kanneganti, 2018; Mantovani et al, 2019). IL-1 family receptors carry the cytoplasmic TIR domain, a shared feature with pathogen sensing TLRs, making them excellent amplifiers of inflammatory signaling (Boraschi et al, 2018). Indeed, nanomolar doses of IL-1α and IL-1β can trigger lethal inflammatory responses in mice and humans (Dinarello, 1996; Lomedico et al, 1984; Smith et al, 1993). Consistently, IL-1α and IL-1β were shown to act as self-amplifying factors and upregulate each other via IL-1R signaling (Dinarello, 2009; Dinarello et al, 1987; Goldbach-Mansky et al, 2006; Greten et al, 2007; Warner et al, 1987). However, studies of IL-1α and IL-1β have produced conflicting results with regard to how these cytokines regulate each other. Studies focused on TLR triggers reported that these self-amplifying positive feedback mechanisms are redundant or not important to amplify the production of IL-1α and IL-1β further (Almog et al, 2015; Copenhaver et al, 2015; Fettelschoss et al, 2011; Glaccum et al, 1997; Labow et al, 1997). These observations differed from studies using a genetic *Il1a* knockout mice (hereafter referred to as *Il1a*-KO^line1^), which showed substantial reduction in IL-1β production when *Il1a* was deleted (Dagvadorj et al, 2021; Gross et al, 2012; Horai et al, 1998), suggesting that IL-1α may regulate IL-1β expression even during TLR activation. These conclusions remained debated and poorly understood for many years.

Therefore, we sought to generate a new line of *Il1a* knockout mice (hereafter referred to as *Il1a-* KO^line2^) using CRISPR-Cas9 technology. The newly generated *Il1a*-KO^line2^ mice showed normal development, with comparable levels of basal immune cells in the blood compared with wild-type (WT) mice. Bone marrow-derived macrophages (BMDMs) prepared from the *Il1a*-KO^line2^ mice showed no defect in expression or activation of inflammasome components in response to PAMPs and live pathogen triggers. Additionally, while the cells from *Il1a*-KO^line1^ showed reduced expression of both IL-1α and IL-1β, *Il1a*-KO^line2^ macrophages had no expression of IL-1α but near-normal expression of IL-1β. Moreover, the *Il1a*-KO^line2^ BMDMs showed a specific requirement of IL-1α for the expression of neutrophil chemoattractant KC/CXCL1, further confirming the functional accuracy of the KO. In summary, we generated and characterized a new line of IL-1α KO mice that improve upon the previous version and have normal IL-1β expression. These mice can be broadly used for future studies on the unique functions of IL-1α and IL-1β to establish their relevance in health and disease and identify new treatment strategies.

## RESULTS

### Generation of the IL-1α KO (*Il1a*-KO^line2^) mouse using CRISPR/Cas9 technology

Although IL-1α has long been recognized as a critical regulator of inflammation and immune responses (Cavalli et al, 2021), its specific functions in physiologic and pathologic inflammatory outcomes in health and disease remain unclear. IL-1α is subjected to complex regulation, and early genetic studies using different knockout mice produced conflicting observations (Dagvadorj et al, 2021; Fettelschoss et al, 2011; Gross et al, 2012; Horai et al, 1998). To clarify the previously observed contradictory roles of IL-1α in IL-1β expression in *Il1a*-KO^line1^ mice, we generated a new line of IL-1α knockout (KO) mice using CRISPR-Cas9 technology, referred to here as *Il1a*-KO^line2^ (Fig. 1). Exons 2-5 of the *Il1a* gene were deleted by using simultaneous injection of two individual gRNAs with human codon optimized Cas9 mRNA (Fig. 1A). We opted to use pronuclear-staged C57BL/6J zygotes for the injections to minimize the background-related genetic issues. Successful generation of the *Il1a*-deficient mice was assessed by targeted deep sequencing and further confirmed by PCR amplification of genomic DNA from the WT and mutant alleles (Fig. 1B), and western blot analysis to confirm the loss of IL-1α protein production (Fig. 2A). Additionally, because IL-1α is a multifaceted cytokine that we postulated may have a role in regulating immune cell phenotypes at basal levels, we evaluated the immune cellularity in the blood from the newly generated CRISPR *Il1a^-/-^* mice (*Il1a*-KO^line2^). We found that these mice did not show any gross abnormalities in the immune cellularity (S. Fig. 1A-B). In sum, we generated a new line of IL-1α knockout mice, *Il1a*-KO^line2^, and confirmed the loss of IL-1α expression with no defects in overall blood immune cellularity.

**Figure 1.**
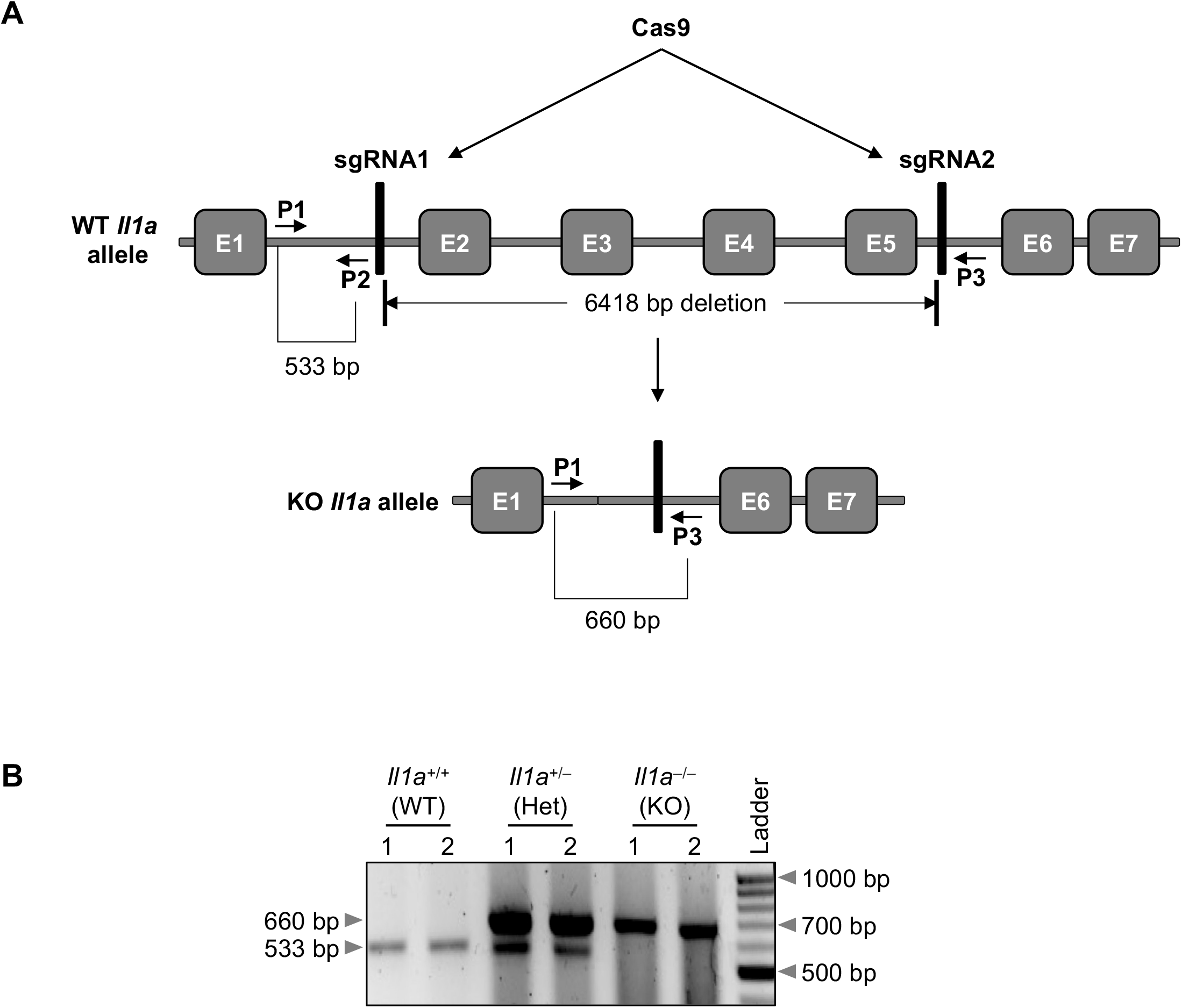
Generation of the *Il1a*^-/-^ (*Il1a*-KO^line2^) mouse using CRISPR/Cas9 technology. **(A)** Two sgRNAs targeting the *Il1a* locus were designed and used to delete exons 2 to 5 (E2 to E5), as described in the materials and methods section. The vertical bars denote the sgRNAs 1 and 2, respectively (depictions are not to the scale) in the genomic sequence. The location of the deleted genomic fragment and the primer-binding locations are depicted using short arrows. (**B**) The PCR amplification of the *Il1a* locus from the DNA of wild-type (WT), heterozygous, (Het), or knockout (KO) mice using the primers (primers P1 and P2 together with P3).

### CRISPR-based genetic deletion of *Il1a* does not affect IL-1β expression or activation

Both IL-1α and IL-1β are known to be highly induced in response to pathogenic insults. Therefore, we next sought to characterize the cytokine expression in cells from the newly generated *Il1a-* KO^line2^ mice in response to PAMPs and pathogens. Treatment of BMDMs with lipopolysaccharide (LPS, a toll-like receptor 4 (TLR4) agonist from Gram-negative bacteria) induced robust and time-dependent expression of IL-1α protein in WT cells but not in *Il1a*-KO^line2^ cells (Fig. 2A). In addition, the induction of IL-1α protein expression was not affected by IL-1β genetic deletion, and the expression of IL-1β in response to LPS was similar in the WT and *Il1a*-KO^line2^ cells (Fig. 2A). In contrast, we observed a delay and reduction in the production of IL-1 β in the previously generated *Il1a*-KO^line1^ cells in response to LPS (S. Fig. 2A).

**Figure 2.**
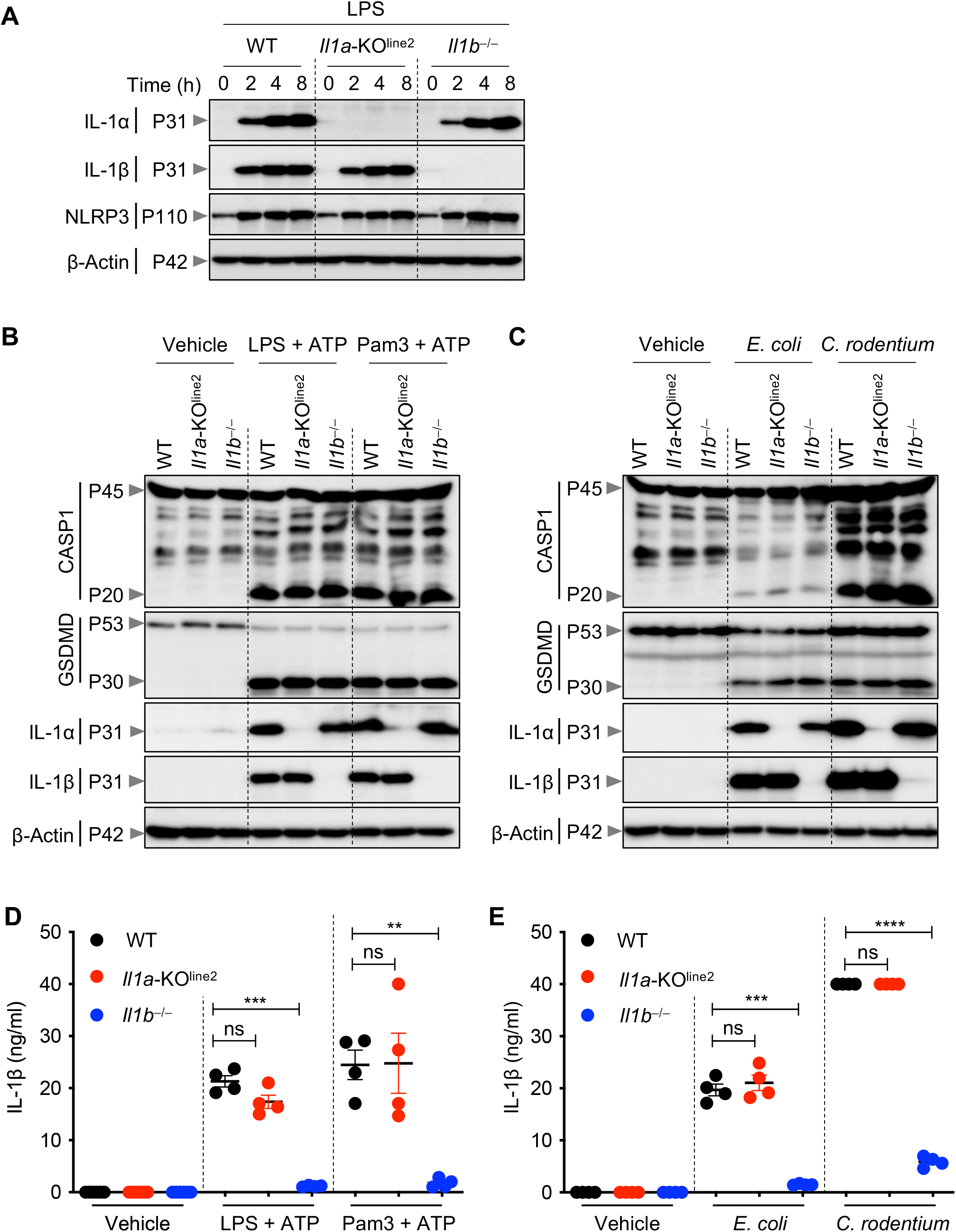
CRISPR-based genetic deletion of *Il1a* (*Il1a*-KO^line2^) does not affect IL-1β expression or activation. (**A**) Western blot analysis of pro-IL-1α (P31), pro-IL-1β (P31), NLRP3 (P110), and β-Actin (P42) in bone marrow-derived macrophages (BMDMs) treated with LPS for indicated times. (**B–C**) Western blot analysis pro-(P45) and activated (P20) caspase-1 (CASP1), pro- (P53) and activated (P30) gasdermin D (GSDMD), pro-IL-1α (P31), pro-IL-1β (P31), and β-Actin (P42) in BMDMs treated with LPS + ATP or Pam3 + ATP for 4 h (**B**), or BMDMs infected with *E. coli* or *C. rodentium* for 24 h (**C**). (**D**–**E**) Measurement of IL-1β release in the cellular supernatants collected from BMDMs treated as detailed in panels (**B**) and (**C**), respectively, for (**D**) and (**E**). Western blot of β-actin was used as loading control. Data are representative of at least two independent experiments (A–E). Data are presented as the mean ± SEM (D and E). Analyses of *the P* values were performed using the *t* test (D and E). ns, non-significant; \*\**P* < 0.01; \*\*\**P* < 0.001; \*\*\*\**P* < 0.0001.

To further understand potential interconnections between IL-1α and IL-1β, we evaluated NLRP3 inflammasome priming, which is known to produce mature IL-1β. We found that IL-1α was not required for upregulation of NLRP3 or IL-1β expression in response to the innate immune triggers LPS, LPS plus ATP, Pam3CSK4 (Pam3) plus ATP, or Gram-negative bacteria *Escherichia coli* or *Citrobacter rodentium* (Fig. 2A–C). Moreover, the activation of canonical and non-canonical inflammasomes, as measured by cleavage of caspase-1 and gasdermin D (GSDMD), were not reduced by deletion IL-1α (Fig. 2B–C). Consistently, IL-1β release was similar in WT and *Il1a-* KO^line2^ BMDMs (Fig. 2D–E). In contrast, using similar experimental approaches, we observed defects in IL-1β expression in macrophages from the earlier *Il1a*-KO^line1^ line, with pronounced reductions in IL-1β expression at early time points in response to LPS, while the induction improved at later timepoints (S. Fig. 2A). We also observed reductions in IL-1β expression in response to NLRP3 inflammasome triggers LPS plus ATP and Pam3 plus ATP (S. Fig. 2B). We did not observe defects in NLRP3 production or caspase-1 and GSDMD activation in *Il1a*-KO^line1^ cells (S. Fig. 2A–B). Together, these results show that while the previously generated *Il1a*-KO^line1^ line had defects in IL-1β production, *Il1a*-KO^line2^ mice did not share these defects.

### CRISPR-based genetic deletion of *Il1a* confirms its critical role in the expression of the chemokine KC (CXCL1)

IL-1α is a pleiotropic cytokine and critical amplifier of inflammation in response to both infection and sterile cellular insults (Cavalli et al, 2021). IL-1α also plays key roles in regulating neutrophil-chemotactic factors such as the chemokine KC (CXCL1) in mice (Gurung et al, 2017). Therefore, to further confirm the IL-1α deletion in the newly generated *Il1a*-KO^line2^ mice and assess its functional effects, we evaluated expression of TNF and KC in response to innate immune triggers. We found that IL-1α specifically was required to produce KC, but not TNF, in response to both PAMP- and pathogen-induced signaling in macrophages; loss of IL-1α resulted in significant decreases in KC release, while loss of IL-1β did not decrease KC release (Fig. 3A–D). Instead, we observed significantly increased levels of KC production in *Il1b^-/-^* cells in response to LPS plus ATP and Pam3 plus ATP treatments (Fig. 3A), suggesting a competition between IL-1α and IL-1β for IL-1R binding in this context, where the increased availability of IL-1R molecules for binding by IL-1α may promote hyper-expression of select inflammatory factors in the absence of IL-1β. Together, these findings confirm the specific role of IL-1α for the release of KC, further supporting the functional relevance of the newly created *Il1a*-KO^line2^ mice for the evaluation of IL-1α— mediated signaling and disease phenotypes.

**Figure 3.**
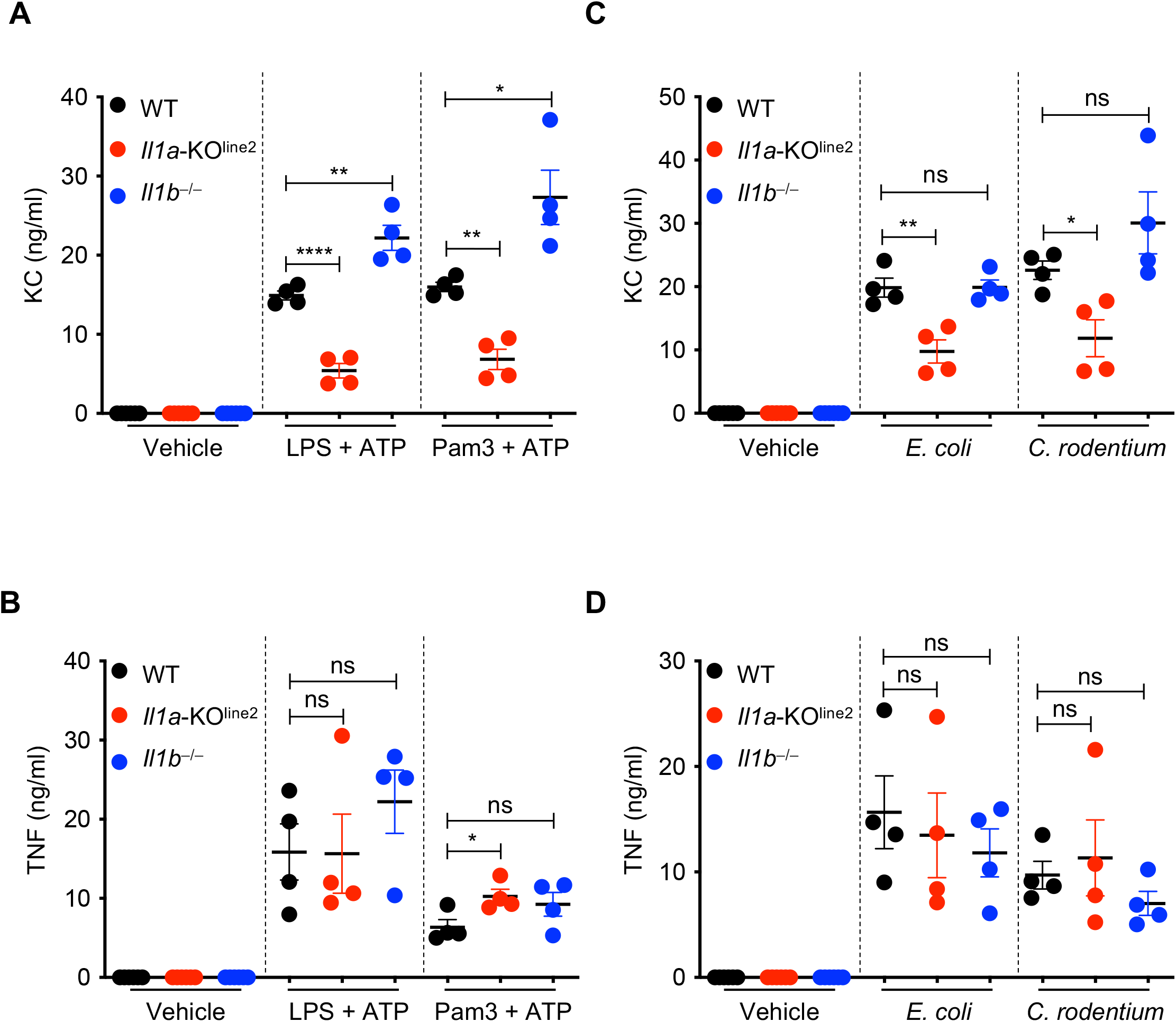
CRISPR-based genetic deletion of *Il1a* (*Il1a*-KO^line2^) confirms its critical role in the expression of the chemokine KC (CXCL1) (**A–D**) Measurement of secreted cytokines KC and TNF in bone marrow-derived macrophages (BMDMs) treated with LPS + ATP or Pam3 + ATP for 4 h (**A–B**) or infected with *E. coli* or *C. rodentium* for 24 h (**C–D**). Data are representative of at least two independent experiments (A–D). Data are presented as the mean ± SEM (A–D). Analyses of *the P* values were performed using the *t* test (A–D). ns, non-significant; \**P* < 0.05; \*\**P* < 0.01; \*\*\*\**P* < 0.0001.

## DISCUSSION

Members of the IL-1 family of cytokines play important roles as inflammatory mediators in host defense but have also been implicated in disease pathogenesis. Therefore, understanding the distinct functions of IL-1 family members is fundamental to our understanding of the molecular basis of disease. Previous genetic models of IL-1α deletion have displayed defects in IL-1β production, making it difficult to determine the distinct roles of these molecules in immune responses. To overcome this obstacle, we report the generation of a genetic deletion of IL-1α in mice using CRISPR technology that did not affect IL-1β induction in response to microbial PAMPs and pathogens. Our findings suggest that expression of IL-1β in response to TLR activation is not affected by loss of IL-1α.

Growing evidence supports that IL-1α and IL-1β have distinct functions (Di Paolo et al, 2009; Eigenbrod et al, 2008; Sakurai et al, 2008). Our findings further confirm that IL-1α is a non-redundant positive regulator of the expression of the chemokine KC in macrophages, which is consistent with earlier studies reporting the specific role of IL-1α in promoting production and recruitment of neutrophils in chronic inflammatory conditions (Gurung et al, 2017; Kono et al, 2010; Lukens et al, 2013; Thornton et al, 2010). However, IL-1β has also been shown to be important for the induction of neutrophil growth- and chemotactic-factors (Cassel et al, 2014; Eislmayr et al, 2022; Gurung et al, 2016; Hsu et al, 2011; Lukens et al, 2014a). Therefore, it is plausible that IL-1β might also contribute to neutrophil-mediated inflammatory conditions as a result of the activation of cell death modalities that drive IL-1β maturation via the activation of caspase-1 or other proteases (Place & Kanneganti, 2019), though this requires further study.

Additionally, previous studies using the earlier *Il1a*-KO^line1^ mice distinguished unique functions of IL-1α and IL-1β in the development of chronic autoinflammatory diseases (Cassel et al, 2014; Lukens et al, 2014a; Lukens et al, 2014b; Lukens et al, 2013; Malik et al, 2016). Our results suggest that the ability to use the *Il1a*-KO^line1^ mice to successfully identify this differential phenotype is due to the chronic nature of the disease. We found that the reduction of IL-1β expression in *Il1a*-KO^line1^ cells was pronounced only at early time points following stimulation, and that prolonged stimulation resulted in similar levels of IL-1β in WT and *Il1a*-KO^line1^ cells, in response to both PAMPs and pathogens. This suggests that WT and *Il1a*-KO^line1^ mice would have similar levels of IL-1β during chronic disease, allowing differential phenotypes between *Il1b^-/-^* and *Il1a^-/-^* mice to be observed.

Given the critical roles of IL-1 family cytokines in inflammation and pathology, these cytokines have been targeted in several therapeutic strategies which have further highlighted unique functions for IL-1α and IL-1β. For example, the recent SAVE-MORE trial showed that anakinra, which blocks both IL-1α and IL-1β, reduced the risk of clinical progression in patients with COVID-19, when co-administered with dexamethasone (Kyriazopoulou et al, 2021). Accordingly, anakinra was authorized for the treatment of COVID-19 in Europe by the EMA. In contrast, the CAN-COVID trial, which was designed to evaluate the efficacy of canakinumab (a specific IL-1β blocking antibody) failed to improve the survival of patients with COVID-19 (Caricchio et al, 2021). These studies further expand the concept that IL-1α plays a dominant and potentially specific role in driving IL-1β-independent inflammatory immune responses and pathology in some contexts.

Together, these observations show that caution should be used when interpreting previous studies and highlight the need to authenticate genetic resources for future work. The development of the *Il1a*-KO^line2^ mouse line, which does not display acute or chronic defects in IL-1β production, may help address many of the critical, long-standing questions in the field regarding the shared and unique functions and context-dependent interdependencies of IL-1α and IL-1β cytokines to improve understanding of the molecular basis of disease and inform therapeutic strategies.

## MATERIALS AND METHODS

### Mice

*Il1b^-/-^* (Shornick et al, 1996) and *Il1a*^-/-^ (*Il1a*-KO^line1^) (Matsuki et al, 2006) mice were both previously described. *Il1a*^-/-^ (*Il1a*-KO^line2^) mice were generated in the current study and are described below. All mice were generated on or extensively backcrossed to the C57/BL6 background. All mice were bred at the Animal Resources Center at St. Jude Children’s Research Hospital and maintained under specific pathogen-free conditions. Mice were maintained with a 12 h light/dark cycle and were fed standard chow. Animal studies were conducted under protocols approved by the St. Jude Children’s Research Hospital committee on the Use and Care of Animals.

### Generation of the new IL-1a KO (*Il1a*-KO^line2^) mouse strain

The new *Il1a*-KO^line2^ mouse was generated using CRISPR/Cas9 technology in collaboration with the St. Jude Transgenic/Gene Knockout Shared Resource facility. Pronuclear-staged C57BL/6J zygotes were injected with human codon-optimized *Cas9* mRNA transcripts (50 ng/μl) combined with two guide RNAs (120 ng/μl each; sgRNA1 for the 5’ of exon 2: AAAAGCTTCTGACGTACCACagg, and sgRNA2 for the 3’ of exon 5: AAGTAACAGCGGAGCGCTT Ttgg (pam sequences are underlined)) to generate a long deletion encompassing exons (E) 2—5 of the *Il1a* gene (Fig. S1A). Zygotes were surgically transplanted into the oviducts of pseudo-pregnant CD1 females, and newborn mice carrying the desired deletion in the *Il1a* allele were identified by PCR agarose gel-electrophoresis (Fig. 1B) and Sanger sequencing. The WT allele was PCR amplified by using the primers IL1a_F1 (5’-GGGCACACGAATTCACACTCACA-3’) and IL1a_R1 (5’-GGAGAACTTGGTTCCTGTTAGGGTGA-3’), and the KO allele was amplified by using IL1a_F1 and IL1a_R2 (5’-TGATTAGCTTCCTTTGGGCTTTGA-3’) primer pairs. The details of the generation of the CRISPR reagents were described previously (Pelletier et al, 2015). The uniqueness of sgRNAs and the off-target sites with fewer than three mismatches were found using the Cas-OFFinder algorithm (Bae et al, 2014).

### Macrophage differentiation and stimulation

BMDMs were prepared as described previously (Gurung et al, 2012). In short, bone marrow cells were cultured in IMDM supplemented with 30% L929 cell-conditioned medium, 10% FBS, 1% nonessential amino acids, and 1% penicillin-streptomycin for 6 days to differentiate into macrophages. On day 6, BMDMs were counted and seeded at 10^6^ cells per well in 12-well culture plates in DMEM containing 10% FBS, 1% nonessential amino acids, and 1% penicillin-streptomycin. iBMDMs (immortalized BMDMs from *Il1a*^-/-^ (*Il1a*-KO^line1^) mice) were maintained in DMEM supplemented with 5% L929 cell-conditioned medium, 10% FBS, 1% nonessential amino acid, and 1% penicillin-streptomycin. Stimulations were performed with LPS alone (100 ng/ml) for the indicated times, LPS (100 ng/ml) or Pam3 (1 μg/ml) for 3.5 h followed by the addition of ATP (5 mM final concentration) for 30 min, or *E. coli* (MOI, 20) or *C. rodentium* (MOI, 20) for 24 h.

### Flow cytometry and analysis of cellularity

The cellular phenotypes of immune cells in the blood were analyzed either by flow cytometry (for T cell subsets and B cells) or by using an automated hematology analyzer machine (for & lymphocytes, % neutrophils, % monocytes, red blood cell (RBC) counts, hemoglobin (HB) quantification, and platelet (PLT) quantification). The following antibodies were used for cell staining: anti-CD19 (APC, clone ID3), anti-CD45.2 (FITC, clone 104), and anti-TCRβ (PECy7, clone H57-597) from Biolegend, and anti-CD8a (efluor450, clone 53-6.7) from eBiosciences. Data were acquired on LSR II Flow Cytometer from BD Biosciences, and analyzed using the FlowJo software (Tree Star), version 10.2 (FlowJo LLC).

### Western blotting

Samples for immunoblotting of caspase-1 were prepared by mixing the cell lysates with culture supernatants (lysis buffer: 5% NP-40 solution in water supplemented with 10 mM DTT and protease inhibitor solution at 1× final concentration); samples for all other protein immunoblotting were prepared without the supernatants in RIPA lysis buffer. Samples were mixed and denatured in loading buffer containing SDS and 100 mM DTT and boiled for 12 min. SDS-PAGE–separated proteins were transferred to PVDF membranes and immunoblotted with primary antibodies against IL-1α (503207, Biolegend), IL-1β (12426, Cell Signaling Technology), caspase-1 (AG-20B-0042; Adipogen), NLRP3 (AG-20B-0014; Adipogen), GSDMD (ab209845, Abcam), and β-Actin (sc-47778 HRP, Santa Cruz), Appropriate horseradish peroxidase (HRP)–conjugated secondary antibodies (anti-Armenian hamster [127-035-099], anti-mouse [315-035-047], and anti-rabbit [111-035-047], Jackson ImmunoResearch Laboratories) were used as described previously (Tweedell et al, 2020). Immunoblot images were acquired on an Amersham Imager using Immobilon^®^ Forte Western HRP Substrate (WBLUF0500, Millipore).

### Cytokine analysis

Cytokines and chemokines were measured by multiplex ELISA (Millipore), as per the manufacturer instructions.

### Statistical analysis

GraphPad Prism 9.0 software was used for data analysis. Data are presented as mean ± SEM. Statistical significance was determined by *t* tests (two-tailed) for two groups.

## DATA AVAILABILITY

The data generated and presented in the current study are provided within the manuscript and the accompanying supplementary figures.

## ACKNOWLEDGMENTS

We thank all the members of the Kanneganti laboratory for their comments and suggestions during the development of this manuscript. We thank R. Tweedell, PhD, and J. Gullett, PhD, for scientific editing and writing support, and Katie Combs and Lauren Kneeland for mouse colony support.

## FUNDING

Work from our laboratory is supported by the US National Institutes of Health (AI101935, AI124346, AI160179, AR056296, and CA253095 to T.-D.K.) and the American Lebanese Syrian Associated Charities (to T.-D.K.). The content is solely the responsibility of the authors and does not necessarily represent the official views of the National Institutes of Health.

## AUTHOR CONTRIBUTIONS

R.K.S.M. and T.-D.K. designed the study. R.K.S.M, R.B., B.K., and B.R.S. performed experiments. A.R.B. and S.P. performed the CRISPR-based knockout generation and initial breeding. R.K.S.M., R.B., and T.-D.K. analyzed the data. R.K.S.M. and R.B. wrote the manuscript with input from all authors. T.-D.K. oversaw the project.

## AUTHOR INFORMATION

T.-D.K. is a consultant for Pfizer. Correspondence and requests for materials should be addressed to.

## SUPPLEMENTAL FIGURE LEGENDS

**Supplemental Figure 1**. **The newly generated CRISPR (*Il1a*-KO^line2^) mice do not show gross abnormalities in blood immune cellularity**

(**A**) Measurement of complete blood counts using automated hematology to analyze percentages of lymphocytes, neutrophils, and monocytes in the total blood cell population, as well as red blood cell (RBC) counts, hemoglobin levels (HB), and platelet (PLT) counts from wild-type (WT), *Il1a^-/-^* (*Il1a*-KO^line2^), and *Il1b*^−/−^ mice. (**B**) Flow cytometry-based quantification of the percent B cells, T cells, and CD4^+^ T and CD8^+^ T cell subsets among the CD45.2^+^ hematopoietic cells from the blood collected from WT, *Il1a^-/-^* (*Il1a*-KO^line2^) and *Il1b^-/-^* mice. Data are representative of at least two independent experiments of 5 to 7 animals per group (A–B). Data are presented as the mean ± SEM (A–B). Analyses of the *P* values were performed using the *t* test (A–B). ns, non-significant.

**Supplemental Figure 2. Previously generated *Il1a^-/-^* (*Il1a*-KO^line1^) cells have defective IL-1β expression**

(**A**) Western blot analysis of pro-IL-1α (P31), pro-IL-1β (P31), NLRP3 (P110), and β-Actin (P42) in immortalized bone marrow-derived macrophages (iBMDMs) treated with LPS for indicated times. (**B**) Western blot analysis of pro-IL-1α (P31), pro-IL-1β (P31), NLRP3 (P110), pro- (P45) and activated (P20) caspase-1 (CASP1), pro- (P53) and activated (P30) GSDMD, and β-Actin (P42) in iBMDMs treated with PBS (Vehi), LPS + ATP, or Pam3 + ATP for 4 h or infected with *E. coli* (EC) or Citrobacter (Citro) for 24 h. Western blot of β-actin was used as loading control. Data are representative of at least two independent experiments (A–B).

