## Supplementary figures and images for "Determining distinct roles of IL-1α through generation of an IL-1α knockout mouse with no defect in IL-1β expression"

### Supplemental Figure 1

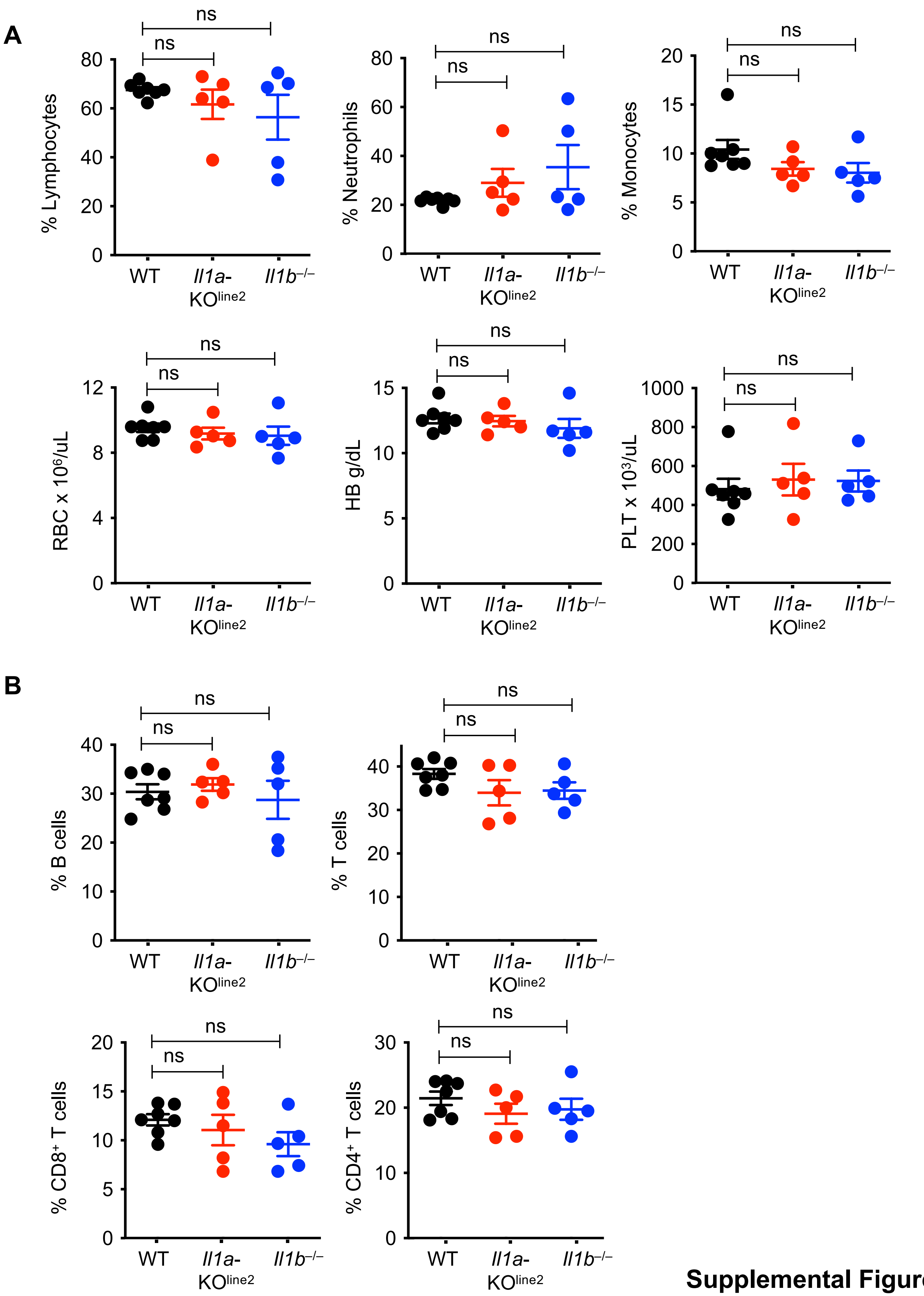


**Supplemental Figure 1**

### Supplemental Figure 2

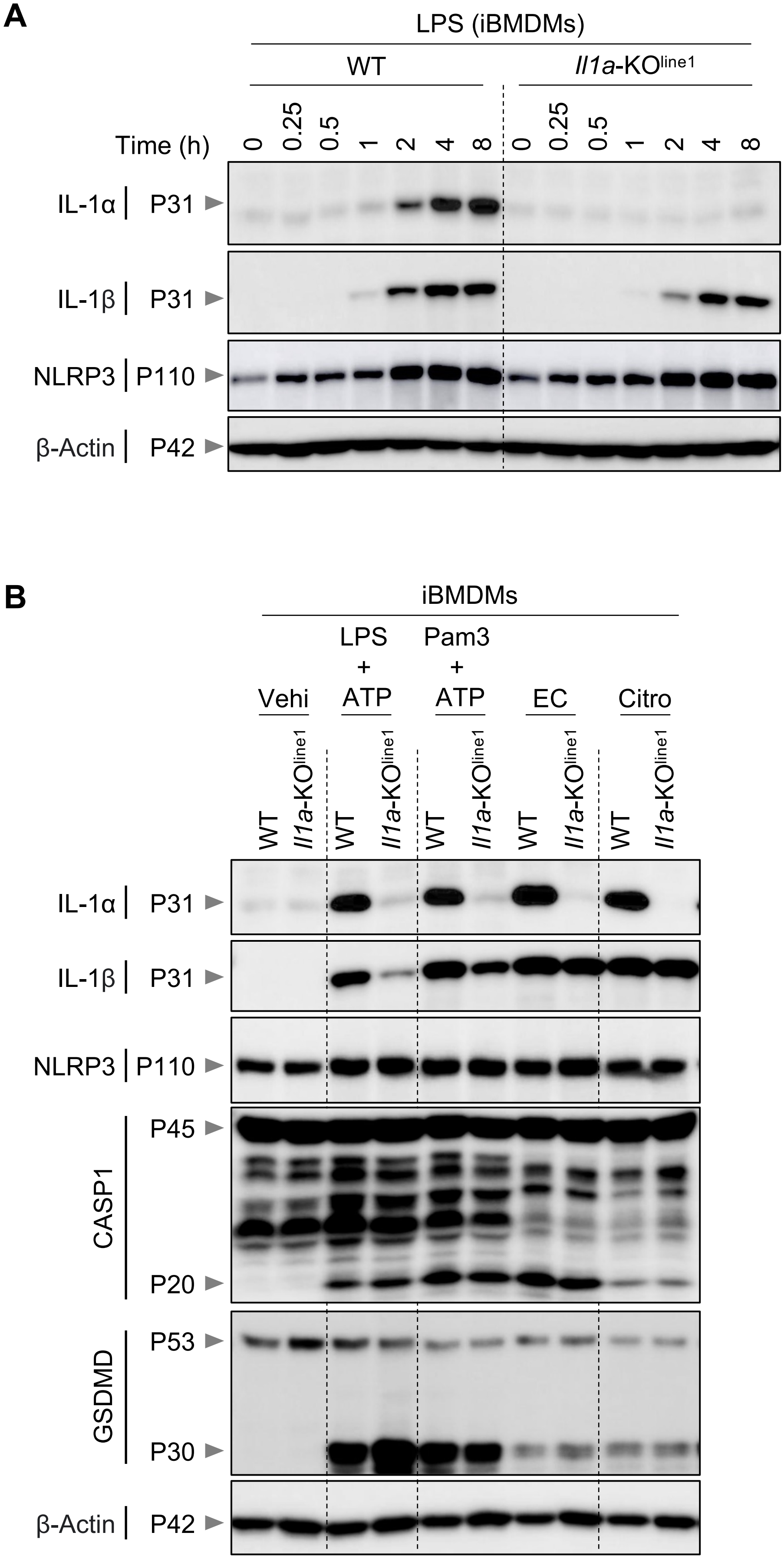


**Supplemental Figure 2**
